# Experimental design for large scale omic studies

**DOI:** 10.1101/532580

**Authors:** Funda Ogut, Jeremy R.B. Newman, Rhonda Bacher, Patrick J. Concannon, Koen J.F. Verhoeven, Lauren M. McIntyre

**Author notes:** these authors contributed equally (Funda Ogut, Jeremy R.B. Newman).

## Abstract

Molecular phenotyping has expanded from small sample sizes to larger complex studies and are now a common element in genetic studies. When large scale studies add a molecular phenotyping component, balancing omics batches for the factors of interest (e.g. treatment), regardless of the initial sample collection strategy always improves power. Where possible, confounding sources of experimental error that are not of interest (sample collection blocks and data collection plates) improves power as does planning batches for molecular phenotyping based on constraints during initial sample collection. Power for testing differences in molecular phenotypes is always higher when accounting for the entire experimental design during modeling. The inclusion of metadata that tracks sources variation is critical to our shared goals of enabling reproducible research.

## Introduction

Experiments using omics technology have transitioned from simple small scale studies to large scale populations. Omics technologies are now deployed in experiments that can encompass genotypes at many loci and/or a diverse array of environmental exposures. Such in depth studies increase our understanding of molecular phenotypes, allowing us to connect molecular signatures to complex traits. In several species, genetic reference panels (GRPs) have been developed and densely genotyped; in human populations large scale studies are now being assayed for omics phenotypes. The attraction to GRPs is the ability to survey the same set of genotypes under different conditions for different phenotypes. GRPs have been used to uncover the genetic basis of flowering time in Maize (1), telomere length (2), male infertility (3), fitness in *C. elegans* (4), and relationships between nutrition and lifespan in *Drosophila* (5). Molecular phenotypes for GRPs have become an important component to understanding complex traits (6), (7), (8) and responses of biological systems to environmental inputs (9, 10). Human genetics has fully embraced both the challenge and the promise of multi-omics experiments with many large scale studies of diseases adding a molecular phenotyping arm.

The application of omics technologies to large experiments brings new challenges to experimental design. For large scale human studies, the wealth of clinical information collected makes repurposing and adding omics data a much more attractive option than a completely new study. For GRPs the ability to compare omics data among treatments/conditions opens and exciting window into complex systems. The goal of experimental design is to maximize the power for detecting and accuracy of estimating effects of interest given practical constraints. In the sample collection phase of large human genetic studies, experimental design considerations are discussed and effects included in subsequent phenotypic models. In crop plants, the fields needed to grow GRPs are large, going beyond what could reasonably be considered a homogeneous environment and a credible randomized complete block design. Yet, there is persistent failure to acknowledge this in the design and analysis of molecular phenotypes. The number of distinct genotypes makes it impossible to collect all study material on the same day, or in a particular circadian window, by a single person, meaning that there will be sources of variation that contribute to the measurement of the molecular phenotype. These are usually factors for which studies of traditional phenotypes would demand a formal experimental design (11). Further complicating the experiments are the data acquisition phases that harbor their own logistical considerations. Thus far how sample collection constraints, should be balanced with design considerations for data acquisition using omics platforms in the subsequent phase of the study has often been neglected in the reporting of molecular phenotypes. Some studies report on their consideration of data acquisition when collecting omics data (16, 17), but most do not provide this level of detail in the reported methods.

Given a specific experimental design for sample collection, is there an optimal way to distribute samples over sequencing lanes, batches, and machines? Experimental design is as critical for acquisition of molecular phenotypes as for sample collection. For example, in RNA-seq, if lanes are confounded with treatments, as suggested by Illumina as a way to minimize the impact of the index hopping (12), inferences about treatment effects will be compromised (13). In metabolomics experiments, effects of batch, machine, and column need to be considered (14). Metabolomics batches are typically small (less than 50) relative to batches for gene expression data (96/384). Effects of initial sample concentration are important to consider for prior to data acquisition to ensure that ionization conditions are comparable across batches (15). Here we use simulations to show, first, how analysis that fails to take into account sources of variation for sample collection and data acquisition results in dramatic reduction of power for inferences, and second; how thoughtful design can be used to optimize power for tests of molecular phenotypes.

Omics experiment have four main stages, sample development, sample collection, sample preparation, and data acquisition (**Fig. 1**). In many experiments, there will be more steps that can introduce variation. Without loss of generality, we consider two general sources of variation (sample development/collection and sample preparation/data acquisition) in order to demonstrate the cascade effects of experimental design choices. The technological or practical capacity for processing data is likely different during sample collection and data acquisition. There are two general randomization approaches for dealing with the transition from sample collection to data acquisition. Batches can be randomized reflecting the original sample collections scheme (conditional randomization – CR) or re-randomized (RR). We evaluate the effect of conditional randomization and re-randomization on the power of tests for genotype and treatment effects. Our findings suggest that power for detection of effects depends upon due consideration of sample collection limitations when planning the data acquisition phase.

**Fig. 1:**
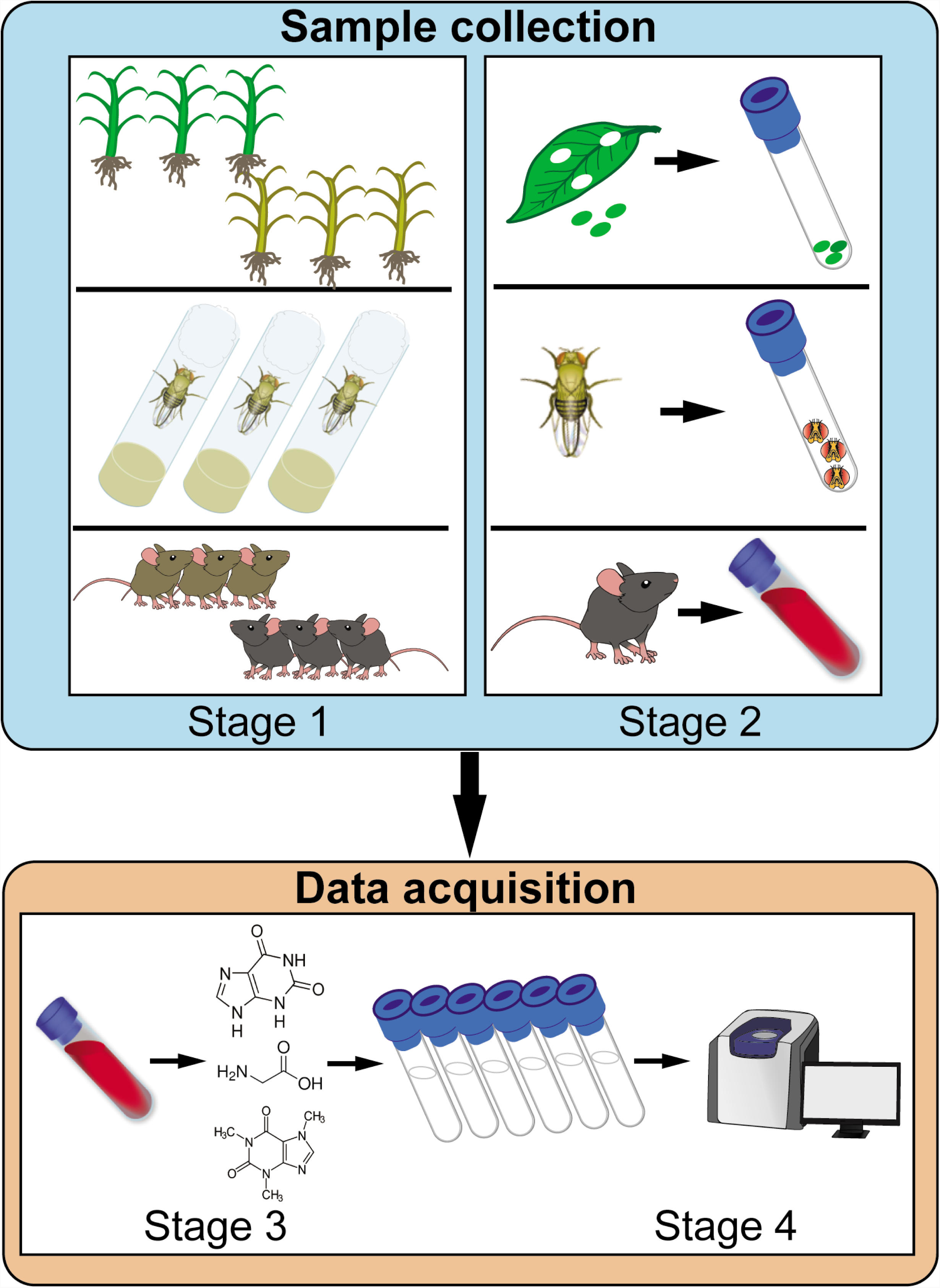
Logistical constraints in the omics experimental process. Stage 1: Sample environment: Nutritional/thermal gradients, treatment applications, cages, vials and other constraints. Stage 2: Sample collection. Circadian rhythm, time from collection to freezing, number of individuals collecting samples. Stage 3: Extraction. The number of extractions that can be simultaneously performed. Stage 4: Data acquisition. Platform, machine variation, reagent variation, index hopping (12) and run time drift (14). Without loss of generality, we can consider Stages 1 and 2 together as sample collection, and Stages 3 and 4 together as data acquisition.

## Results

Molecular studies often have broad hypothesis (for example: there is a difference in gene expression between the control and treatment for at least one gene; or changes in at least one of the measured metabolites reflects the underlying metabolic cause of disease and will be independently predictive of condition in other samples) and reuse of the data for other hypotheses is common. As different hypotheses will suggest different designs to optimize power, it is helpful to define primary and secondary goals for the study before choosing a design. However, when such studies are added later to an existing study, this is not always possible. Here we consider the impact of the sampling design on two common null hypotheses for molecular phenotypes: the absence of genotypic differences and the absence of differences in treatment effect between genotypes. Practical advice for omics data acquisition is provided for situations where there is no ability to integrate the design for sample collection and omics data acquisition.

### Tracking the experimental design is crucial

Although advanced methods exist to account for hidden sources of variation (e.g., 17), the most common approach to statistical testing in molecular phenotypes is to ignore the experimental design in the sample collection phase and sometimes also during the data acquisition. While normalization for data acquisition batch effects is common, when batch and treatment effects have complicated relationships normalization strategies are likely to fail (17) and can lead to radically misleading results (18). We compared models that ignore block effects (during sample collection) and batch effects (during omics data acquisition) to models that account for these effects. The models that do not track the design in either or both stages of the experiment have less power for detecting genotype effects than those that do track the design (**Fig. 2**). Models that ignore block or batch effects, even when they are smaller than the biological variation, have less power to detect effects of interest than when variance between blocks and batches is relatively large and accounted for in the model. Other trends, such as decreasing power when batch variance increases are intuitive, yet striking in the magnitude of the impact.

**Figure 2:**
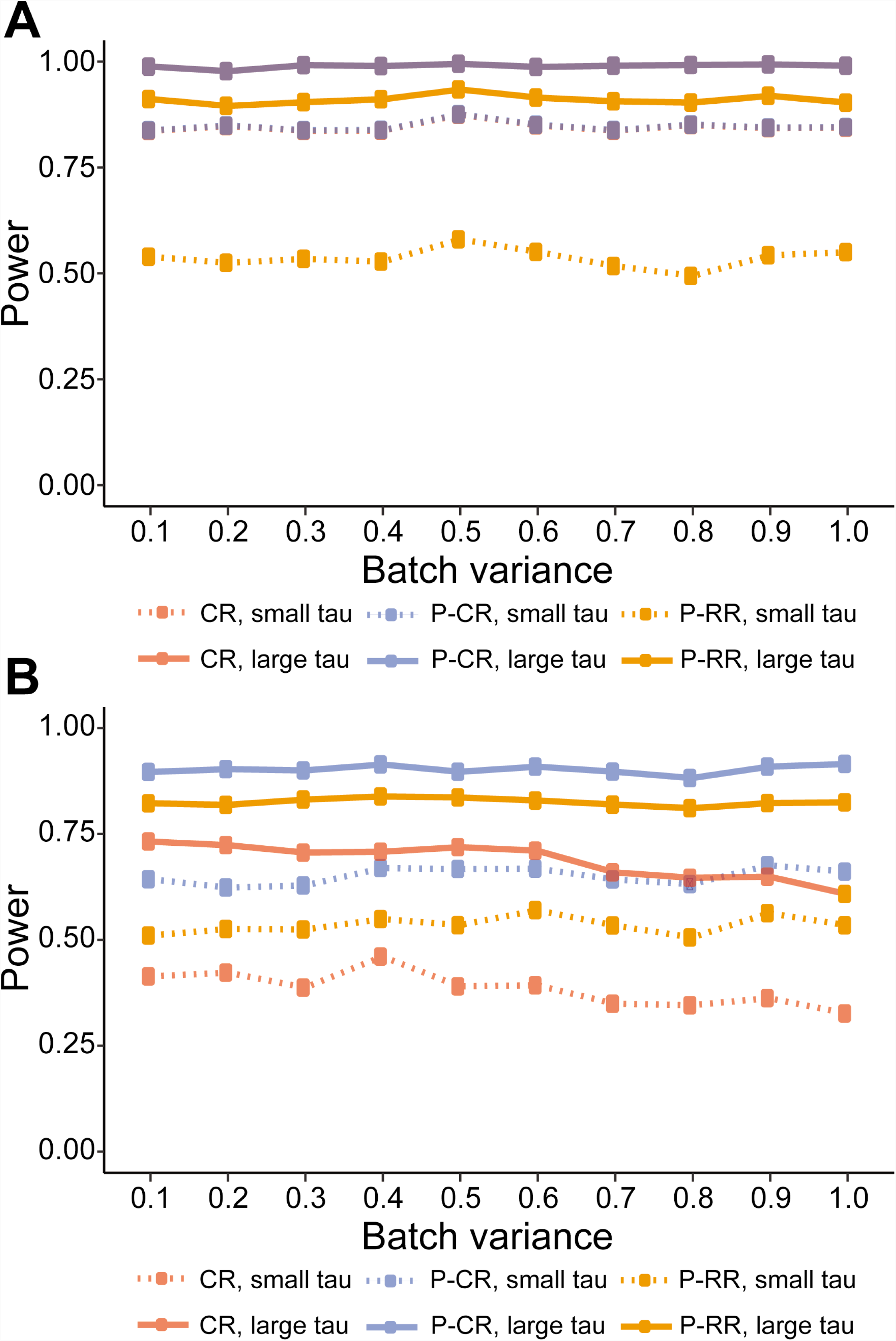
Models that reflect the design have higher power to detect genotypic differences. Including the block/batch effects in the model is crucial to testing for the genetic effect (QTL effect size = 1.2, batch variance is large, replicate variance = 32, error variance = 15). (**A)** When the block size is bigger than the batch size and (**B)** when block size is less than batch size. Failing to incorporate block and/or batch effects has a large effect on power. Accounting for batch effects has higher power than failing to account for batch effects and the additional impact of accounting for block are large with power gains dependent on the size of the block with high block variance increasing the importance of accounting for the block effect.

### Conditional randomization is better than re-randomization in tests for genotype effects

How to best match samples between sample collection blocks and data acquisition batches? One option is to keep the experimental units (e.g. genotypes) collected in the same sample collection block together in the same batch for data acquisition (conditional randomization, CR). Alternatively, the samples can be randomized to new batches for data acquisition without regard to the original sample collection block (re-randomization, RR) (**Fig. 3A**). For illustration consider the simplest case where all samples for a single replicate can be collected as a single block and then data acquired in batches of equal size to the blocks. Here, CR improves power for detecting genotypic differences (**Fig. 3B**). When the experimental errors of blocks and batches are completely confounded they are jointly estimated as a single effect leaving more degrees of freedom for the estimation of the residual error (See supplementary material for a theoretical argument). Effectively, all genotypes are measured simultaneously in the same ‘environment’, thereby reducing the variance in the between genotype comparison. In contrast, complete re-randomization (RR) results in the need to separately estimate block and batch effects. Effectively constructing different environments for each genotype and making the accurate estimation of these effects necessary for comparing genotypes. In a GRP, the small number of replicates relative to the number of genotypes limits the estimation of design effects such as block and batch even in this relatively simple cases where the block and batch variation are independent from each other and from the effect of genotype (See Supplement). In more realistic scenarios where sampling is conducted in partial blocks within replicates (**Fig. 3C**), CR can have much higher power for the detection of genotype effects than RR. (**Fig. 3D**). In comparisons of genotypic effects CR is markedly more powerful than RR when comparing genotypes in the same partial block/batch combination. When comparing genotypes across partial blocks/batches power is comparable in CR and RR (**Fig. 3D**). There is then nothing to be lost, and some power to be gained by using CR designs for data acquisition of molecular phenotypes in the comparison among genotypes.

**Fig. 3:**
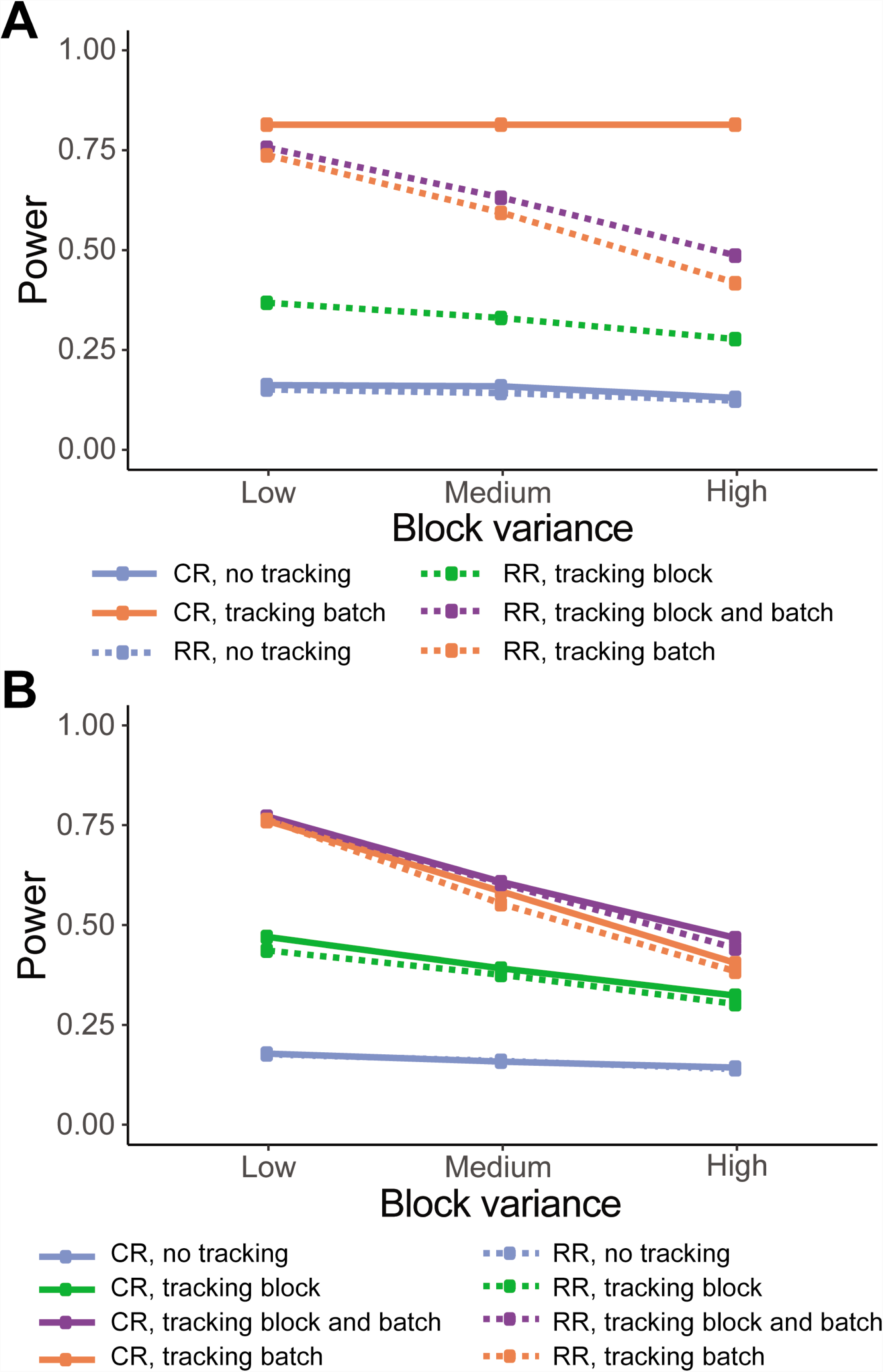
Conditional Randomization versus Re-Randomization for tests of genotypic effect. (**A)** Each full replicate contains all genotypes and is processed simultaneously. For data acquisition, CR and RR are compared. (**B)** A QTL simulation with a single main effect for 96 recombinant inbred lines and four replicates is considered. Power for the test of the QTL effect is shown on the y-axis when a single replicate has a single block (n=96) with moderate variance across replicates. Batch variance is plotted along the x-axis as a function of the replicate/block variance (1/3x – 2x). Two effect sizes for the QTL are shown. (see **Supplementary Table 1**). Power for detection of a difference is estimated as the number of rejections at a nominal p-value of 0.05 out of 1,000 iterations for the test of null g_j_=g_j’_ using the model Y_ij_= replicate_i_+genotype_j_ + error_ij_ for CR and Y_ijk_=replicate_i_+batch_k_+genotype_j_+ error _ijk_ for RR. Batch is not included in the CR model as it is completely confounded with replicate. (**C)** Partial blocks consisting of a subset of genotypes. Here CR keeps genotypes from a block together in the batch for data acquisition and RR ignores the block when randomizing to batches. Randomizations between replicates are independent. (**D)** A natural population simulation (See **Supplementary Table 3**). On the y-axis is power calculated as in B and on the x-axis is the variance due to batch, with a block variance of 1 and error variance of 0.4. As the variance between batches increases the power for the test of the genotype effect decreases in RR. Power between genotypes in different batches in CR is virtually identical to the RR design, while power between genotypes in the same batch in the CR design is much higher. For the P-RR design the average power for genotype effects is shown without regard to the batch.

### The relative size of the block and batch matters

For some experimental systems (e.g. maize) the sample collection phase can have relatively large blocks, while in other systems (e.g. mouse) the sample collection phase is typically constrained to much smaller blocks. In human genetics sample collection is a process that is reflective of the unique constraints of these studies. The relative size of the sample collection block compared to the data acquisition batch varies among studies. While gene expression assays can be processed in relatively large batches (RNA-seq in multiplexed batches of 96/384), smaller batch sizes are more common for other omics platforms (12-48 in metabolomics). The impacts of batches can be easily demonstrated (13, 15)(**Supplementary Fig. S3 and S4**) The size of the batch correspondingly affects the number of batches needed, with smaller batch sizes requiring a larger number of batches to complete an experiment. We evaluated power to detect a QTL effect given different block and batch size/number. We compared RR and CR for different amounts of variance (**Fig. 4, Supplementary Fig. S2**). Power is similar when the number of blocks is small relative to the number of batches (**Fig. 4a**). For medium and high variance in blocks and batches CR has higher power. Power is higher when the size of the block is equal to or greater than the size of the batch. (**Fig. 4a**) In CR, the highest power is always when block and batch size match (**Fig. 4b**). In contrast, for RR larger blocks are more important than larger batches (**Fig. 4b**). When block variance is low block size does not matter (**Fig. 4B**). When block and batch variance are equal and the variance is low or high CR and RR are very similar in their power to detect QTL effect (**Fig 4c**). As the block and batch size increases, CR improves in power faster than RR (**Fig 4c**).

**Fig. 4:**
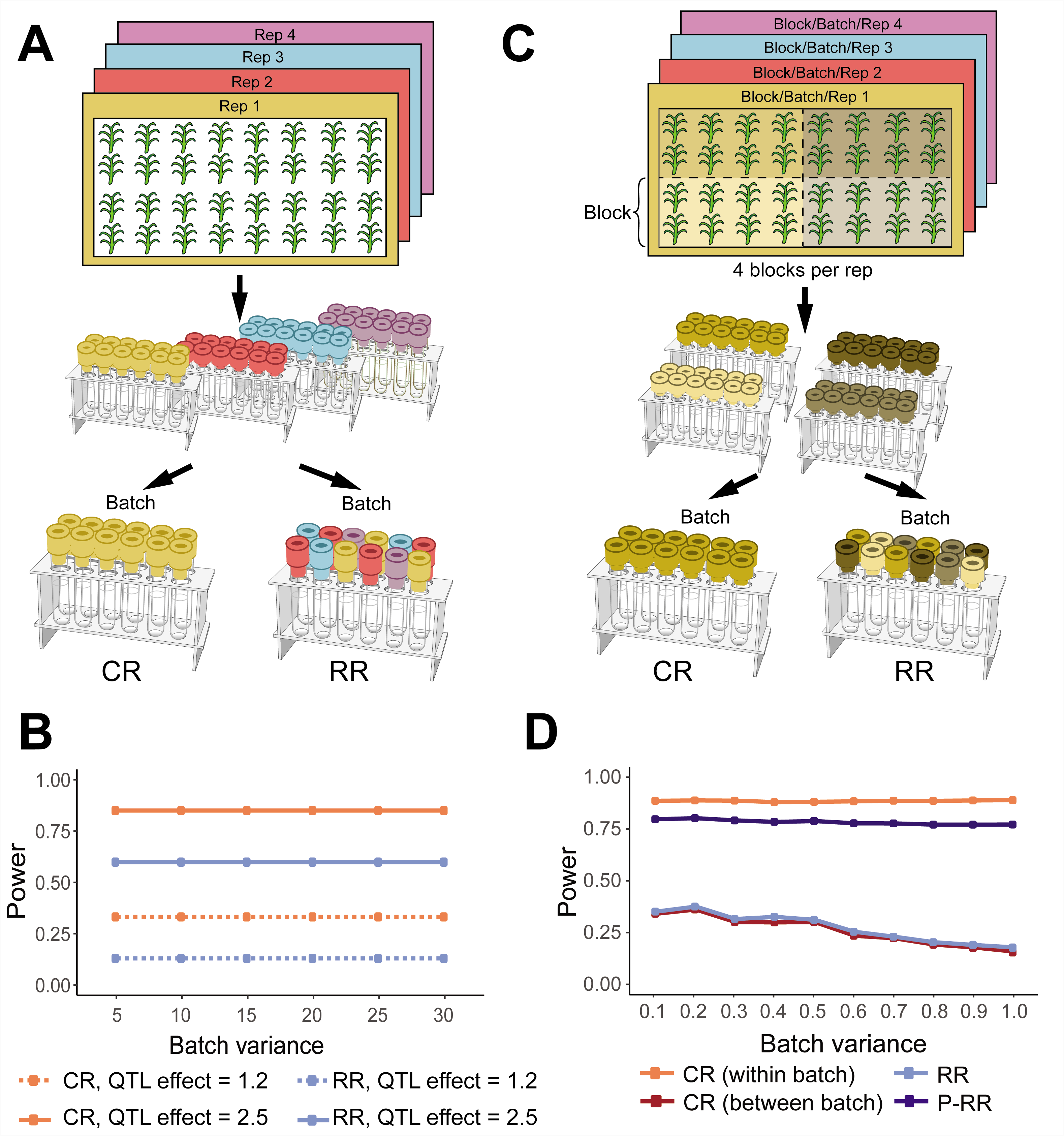
Effect of block and batch size on power. (**A**) A simulation was conducted for a QTL with a single main effect for 192 RILs and 4 reps, with a QTL effect of 1.5, replicate variance of 32 and error variance of 15. Block and Batch variance are equal. The ratio of block size/batch size on the X axis power on the Y axis. Full simulation conditions given in **Supplementary Table 2** (**B**) Effect of batch and block size on power when batch and block variance are high (variance=24) and low (variance=8), with QTL effect of 1.2, replicate variance of 32 and error variance of 15. A composite when batch size is 96 is also shown. When variance is high (variance = 24), the power is higher when batch sizes are smaller than block sizes, and variance has little impact on power for the CR model. The RR model has less power than the CR, irrespective of batch and block size. For both models, when block variance is low block size does not matter. Full simulation conditions given in **Supplementary Table 3**.

### Power for testing treatment effects depends on pairing treatments

The effect of treatment, such as tissue differences in mice, drought or other environmental stress effects in plants, and sex differences in Drosophila may be sampled in the same block for a particular genotype (incomplete block design, IBD; **Supplementary Fig. S1A**) or may require separation for sampling (e.g. planting density (19)) (split-plot design, SPD; **Supplementary Fig. S1B**). The SPD has lower power for the whole plot (treatment) effect relative to the IBD for phenotypes sampled during the sample collecting phase (11). Most human genetic studies collected over multiple sites will be some variant of the SPD as sites are often different for confounding factors such as age and race. We simulated 192 genotypes in a natural population with varying treatment effects and without interactions between the treatment and genotype (**Supplementary Table 3**) in both an IBD and SPD. Data acquisition was simulated in three ways: 1. in batches that retain the block randomization (CR); 2. pairing samples from the same genotype and block but from different treatment levels (for instance treatment-control) and then randomizing these pairs into batches (P-RR) and pairing samples from the same genotype and block but from different treatment levels (for instance treatment-control) and then keeping half of the genotype treatment pairs together in a batch (P-CR) (**Supplementary Fig. S1A,B**). Power is lower for SPD than IBD for the test of treatment, since the treatments are confounded with block (**Fig. 5**). However, as long as the treatments for a genotype are paired in the data acquisition stage (P-RR and P-CR), power in the SPD is similar (**Fig 5**). The implication for human genetic studies is that molecular phenotypes should be acquired in batches balanced for tests of the effects of interest, regardless of the initial sample collection.

**Fig. 5:**
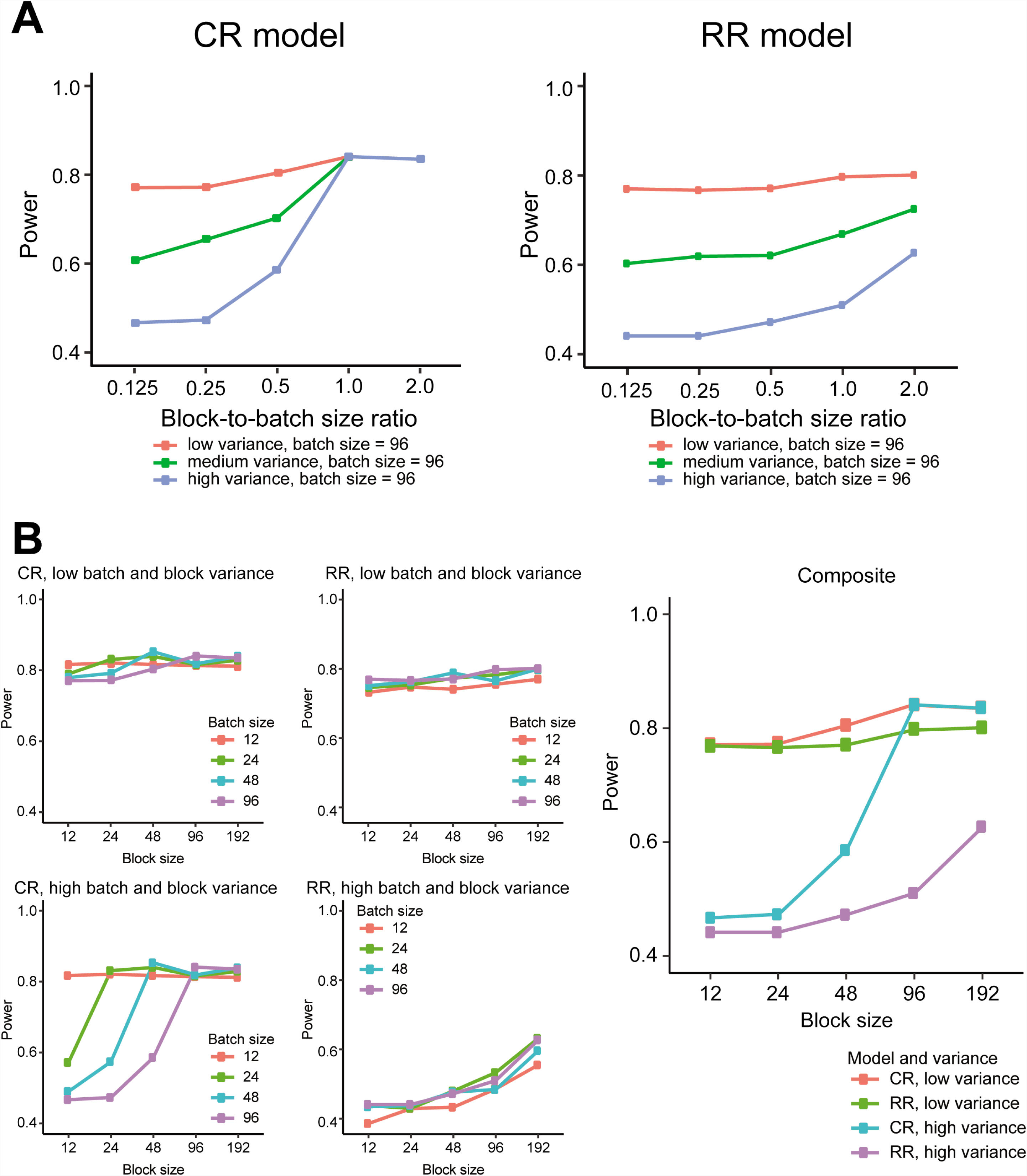
Power of the test of treatment in IBD and SPD with different batch randomization strategies. A simulation was conducted for a natural population (**Supplementary Tables 3/4 and Fig. S1**). Batch variance is on the X axis and power on the Y axis. Block variance is 1, and treatment variance is 1. (**A**) The IBD design RR has less power than the CR. In this comparison the treatment effects are paired at both sample collection and for data acquisition in CR while treatments are randomized in the block and again in the batch, some pairing will happen by chance. Note that the CR (red lines) and paired-CR (P-CR; blue lines) designs have the same result. (B) The SPD design. Here treatments are in different blocks by necessity. CR keeps treatments separated and confounds treatment with batch. P-CR pairs treatments and keeps as much of the block structure as possible in batches and P-RR pairs treatments but randomizes pairs with respect to batches.

## Discussion

One of the barriers to successful omics data analysis and subsequent integration is the treatment of experimental effects as a post-hoc normalization problem. By focusing on *post-hoc* removal of technological effects, strong assumptions are needed. *Post-hoc* normalization strategies are not guaranteed to succeed (18), and they cannot fix issues with confounded effects in the design (11, 13). Tracking the experimental design and including those effects in the model when testing for the effects of interest improves power dramatically. In large studies of phenotypes such as plant yield or bone density, it is common to account for sources of variation in the design and in particular at the time of measurement. While tissue collection and downstream data acquisition for molecular phenotypes may be more complex than organismal phenotypes, the increased cost and complexity of phenotyping does not change the necessity for careful experimental design. Studies likely consider these elements during planning, but fail to report these details. More exposition of sample collection protocols in and the design of the data collection strategies are critical to improving reproducibility. Further, fitting models that explicitly account for the complete design provide substantial gains in power compared to ignoring these effects. Randomization protocols for omics data that only consider the logistical constraints for omics data acquisition are not necessarily optimal designs. Optimal strategies for linking design between sample collection and data acquisition depend on logistical constraints (e.g. the relative size of blocks and batches) but in all cases power gains can be achieved by accounting for the experimental design for the entire scope of the experiment.

Optimal designs will depend upon the specific hypotheses to be tested. For a hypothesis about a treatment effect, if the sample collection forced treatments to be in separate environments (a “split plot”) then a randomization for sample collection that pairs the same genotype with all treatments for data acquisition will have more power than keeping the treatments separated in batches. In human genetic studies of molecular phenotypes, it is common to account for covariates (e.g. age, sex) as well as technical factors such as batch in the statistical model (17, 21). Human genetic studies are often “split plot” designs with some confounding of sample collection with factors of interest/importance like sex and age, treatment arm, etc. In these cases, we recommend a restricted re-randomization that balances potentially important covariates over batches, rather than an unrestricted re-randomization (**Fig 5, Supplementary Fig. 1**).

When studies are repurposed for omics experiments, it is of critical importance to both keep track of sources of variation in the original study, and to balance omics blocks for the treatments of interest for the omics experiment. Similar to (13) our simulations show that for comparisons between conditions pairing samples by condition/treatment during omics data acquisition minimizes the confounding between technical sources of variation during omics data collection and the hypothesis of interest. Treatments should not be confounded with sources of variation during data acquisition, e.g. lanes, despite issues that may arise (e.g index hopping). Instead in Illumina studies larger multiplexes, and more technical replicates are preferred. The use of unique i5/i7 barcode combinations also facilitates the detection of this phenomenon. For metabolomics, balancing the small batches so that important contrasts among samples are captured within the batch is critically important (e.g. (15)).

The challenge to large scale molecular phenotyping studies, particularly those investigating multi-omics, is in the number and scope of hypotheses. Secondary data analyses are an important part of the justification for conducting these experiments. Logistical constraints create situations where it is unlikely that the optimal design for all possible hypotheses can be developed – there will be trade-offs. While difficult, investigators should actively manage these trade-offs during design of the experiments. It is of critical importance that the experimental design process is realistic about the sources of variation in the entire experimental process. Ignoring sources of variation can result in dramatic loss of power. For investigators partnering with companies for data acquisition services, information about batches, and randomization within and between batches is of critical importance. Companies that do not disclose this information may confound sources variation obscuring true differences and potentially leading to false positives due to technological effects. Lack of information about batches will lead to increased challenges in reproducing results.

The power of large molecular studies can be improved by jointly planning the experimental design of data acquisition and sample collection to optimize block and batch sizes. We show that higher power is obtained, with the same data, if the statistical model of molecular omics phenotypes explicitly includes the experimental design of both the sample collection and the data acquisition phases – and that further power gains are possible if the design for data acquisition is reflective of constraints of the sample collection phase. If batch variance is expected to be high, smaller batches with samples handled jointly throughout the process will maximize power. Researchers should avoid the temptation to continually re-randomize samples during omics data processing. Instead, a randomization that balance for effects of interest that is maintained will both result in fewer handling errors by technical staff, and higher power for hypothesis tests.

Even when the technical variance is small relative to the biological variation, substantial power can be gained by accounting for technical sources of variation. The immediate implication is that failure to share meta-data related to sample collection and data acquisition, will result in increased difficulty in reproducing results. The important endeavor of increasing the transparency, reliability and reproducibility of results demands that detailed information about how data are collected, in all the stages of the experiment, are critical to future studies.

## Methods

### QTL simulation

Data were simulated for 96 inbred lines (RILs) with one major QTL and 4 independent replicates according to the model ***Y***_***kijl***_ = ***u*** + ***R***_***k***_ + ***G***_***i***_ + ***ζ***_***ki***_ + ***P***_***j***_ + ***γ***_(***jl***)_, Y_kijl_ trait value of genotype *i*^*th*^ in rep *k*^*th*^, u, population mean u = 0, R_k_ replicate effect 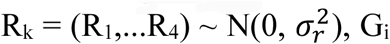 genotype effect G_i_ = (G_1_,…,G_96_) ∼ N(α1,σ^2^) where the additive genetic effect was simulated with a QTL effect (α_i_) using R/qtl (20). Independent of the QTL we simulated random variation from sample collection 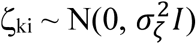, data acquisition 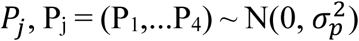, and random variation due to noise 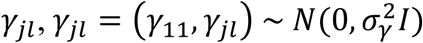 where random noise was scaled relative to 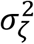. By varying α, 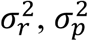 and 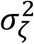 a total of 180 conditions were simulated with 1000 iterations each (**Supplementary Table 1**). The different randomization elements of the design were evaluated by arranging the block/batch variance according to the design being evaluated.

**Table 1:** Simulation conditions for the experiment to test RR and CR in the basic design where a single replicate can be placed in a single block during sample collection. All combinations below were simulated for a total of 180 conditions with 1000 iterations each. Each simulation had four replicates of 96 genotypes for a total of 384 experimental units. Values of 15 were considered moderate variance.

| QTL effect size ($\alpha_g - \alpha_{g'}$) | $\sigma_g^2$ | $\sigma_{e1}^2$ | $\sigma_r^2$ | $\sigma_p^2$ |
| --- | --- | --- | --- | --- |
| 1.2, 1.5, 1.8, 2.0, 2.5 | 1 | 15 | 5, 10, 15, 20, 25, 30 | 5, 10, 15, 20, 25, 30 |

**Table 2:** Simulation conditions for the experiment to test RR and CR when there is blocking with varying batch and block sizes. All combinations below were simulated for a total of 360 conditions with 1000 iterations each.

| QTL effect<br>size ($\alpha_g - \alpha_{g'}$) | $\sigma_g^2$ | $\sigma_{e1}^2$ | $\sigma_r^2$ | $\sigma_f^2$ | $\sigma_p^2$ | Block ($f$) size | Batch ($p$)<br>size |
| --- | --- | --- | --- | --- | --- | --- | --- |
| -1.2, -2.5 | 3 | 15 | 32 | 8,16,24 | 8,16,24 | 12,24,48,96,192 | 12,24,48,96 |

**Table 3:** Simulation conditions for the experiment to test RR and CR in the basic design for where a single replicate can be placed in a single block during sample collection. All combinations below were simulated for a total of 10 conditions with 50 iterations each with four replicates of 192 genotypes for a total of 768 experimental units.

| Genotype<br>mean ($\mu_g$) | $\sigma_g^2$ | $\sigma_{e1}^2$ | $\sigma_r^2$ | $\sigma_p^2$ |
| --- | --- | --- | --- | --- |
| 1-192 | 0.2 | 0.4 | 1 | 0.1,0.2,0.3,0.4,0.5,0.6,0.7,0.8,0.9,1 |

**Table 4:** Simulation conditions for the experiment to test RR and CR in the basic design where a single replicate can be placed in a single block during sample collection. All combinations below were simulated for a total of 180 conditions with 1000 iterations each with four replicates of 96 genotypes for a total of 384 experimental units.

| Genotype<br>effect<br>($\mu_g$) | $\sigma_g^2$ | treatment<br>effect<br>($\mu_t$) | $\sigma_t^2$ | $\sigma_{e1}^2$ | $\sigma_r^2$ | $\sigma_b^2$ | $\sigma_p^2$ |
| --- | --- | --- | --- | --- | --- | --- | --- |
| 1-192 | 0.2 | 2 | 0.5, 1, 2 | 0.4 | 0.7 | 0.1,0.2,0.3,0.4,0.5,0.6,0.7, 0.8,0.9,1 | 0.1,0.2,0.3,0.4,0.5,0.6,0.7, 0.8,0.9,1 |
| 1-192 | 0.2 | 3 | 0.5, 1, 2 | 0.4 | 0.7 | 0.1,0.2,0.3,0.4,0.5,0.6,0.7, 0.8,0.9,1 | 0.1,0.2,0.3,0.4,0.5,0.6,0.7, 0.8,0.9,1 |
| 1-192 | 0.2 | 10 | 0.5, 1, 2 | 0.4 | 0.7 | 0.1,0.2,0.3,0.4,0.5,0.6,0.7, 0.8,0.9,1 | 0.1,0.2,0.3,0.4,0.5,0.6,0.7, 0.8,0.9,1 |

### Population simulation

#### Block design

Datasets were simulated for 192 genotypes with 2 treatments and 4 replicates. For each replicate, b blocks were simulated from multivariate normal distribution (MVN) assuming independent block effects. For each replicate, genotype/treatment pairs were assigned to each block randomly (note that genotypes within blocks vary rep to rep – genotype set in rep1, block1 are not the same genotype set in rep 2, 3 or 4, block 1). For each simulated dataset, the phenotype was simulated as the sum of genotype effect, treatment effect, rep effect, block effect and random error for i = 1 to 192 genotypes, j = 1 to 2 treatments (control vs treatment), k = 1 to 4 replicates and l = 1 to b blocks.

#### Split Plot Design

Using the same parameters as described above for the genotypic effects. For each replicate, b blocks were simulated from multivariate normal distribution (MVN) assuming independent block effects. For each replicate, 4 of the blocks were randomly assigned to treatment 1 and the other four of them assigned to treatment 2 and 192 genotypes were assigned randomly (note that genotypes within blocks vary rep to rep – genotype set (48 genotypes) in rep1, block1 are not the same genotype set in rep2, 3 or 4, block 1). For each simulated dataset, the phenotype was simulated as the sum of genotype effect, treatment effect, rep effect, block effect and random error for i = 1 to 192 genotypes, j = 1 to 2 treatments (control vs treatment), k = 1 to 4 replicates and l = 1 to n blocks.

## Acknowledgements

National Institute of Health (GM128193, U2CES030167, DP3DK085678), NSF PGRP 1238030, Bayer Traits in Seeds Grant Program. Gary Churchill, James Holland, Dirk-Jan de Koning and Bruce Weir have all provided helpful discussion that has influenced the direction of this work.

## Author Contributions

FO and JRBN contributed equally to this work. FO was responsible for the majority of the coding and parameter selection in the simulations while JRBN designed and constructed all of the figures. FO and JRBN participated in simulation and figure development and both contributed to the writing of the manuscript. RB developed the theoretical expectations and contributed to the writing of the manuscript. PC provided insight into the impacts in human genetics, provided the human data summaries of technical effects in the supplement, and contributed to the writing. KJFV contributed to the development of the hypotheses, design of simulations, and the writing of the manuscript. LMM initially posed the problem, developed the hypotheses to be tested, contributed to the design of the simulations, the summary of the results and had major responsibility for the writing of the manuscript.

## DESIGN-OMICs simulations

Using these indexes throughout

Replicate r 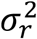

Block f 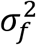

Batch p 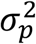

Genotype g 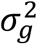

Individual i

Treatment t

### QTL based simulations

#### Simulation 1. Conditional Randomization vs Re-Randomization across replicates

For the first simulation, our goal was to understand the effect of variation due to sample collection (block) and variation due to data acquisition (batch). In this simple situation a single replicate fits into a single block for sample collection and a single batch for data acquisition. Data were simulated for 96 inbred lines (RILs) with one major QTL in 4 independent replicates (4 blocks each containing 96 genotypes). After sampling, experimental units were either kept together on a plate for data acquisition (CR) or randomized without regard to the initial sample collection phase (RR).

We simulated each source of variation independently. Here block effect is the effect of replicate. For *r* = 1 to *r* replicates, the effect of each replicate is drawn independently from 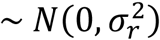. For *p* = 1 to *p* batches, the effect of each batch is drawn from 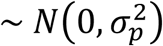. For *g* genotypes we simulated a single QTL with an additive effect using R/qtl (Broman et al. 2003) and drew from ∼ *N*(*α*_*g*_ - *α*_*g*′_, 1), where *α*_*g*_ - *α*_*g*′_ was the QTL effect. Each individual also had two sources of random error associated with each observation, one at sample collection 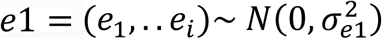 and one at data acquisition 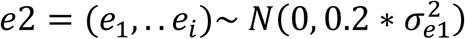.

The trait value for an individual *i* in rep *r* and batch *p* is calculated as:

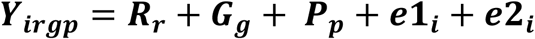

In general, let *n*_*g*_ be the set of individuals having genotype g, and |***n***_***g***_| be the number of individuals in *n*_*g*_, similarly 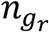 and 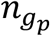 will denote the set of replicates and batches for individuals having genotype g. The expected mean trait value for genotype g is (without loss of generality, the expectation is conditional on the random effects rep and batch):

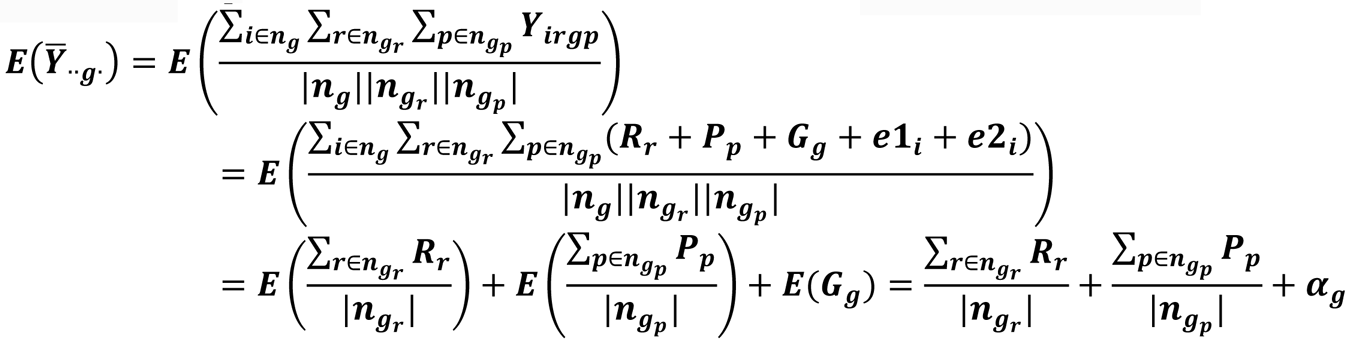

The generalized least squares estimator of the QTL effect between genotypes g and g’ is:

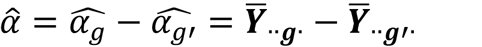

In CR, 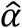 is unbiased as shown below since the replicate and batch effect are equivalent between the two genotypes means 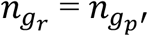 for all *r* and *p*.

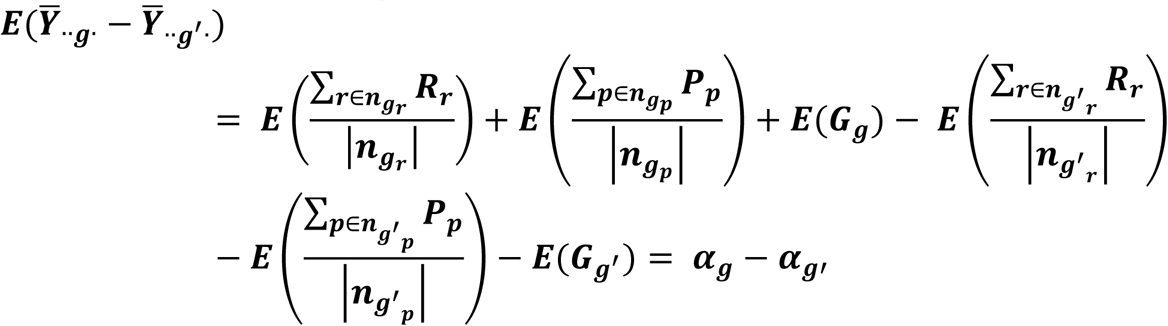

In RR, genotypes are initially randomized for the replicates in the same way, where each genotype occurs once in each replicate. The difference is that during batch assignment there is no consideration of the replicate that the sample came from. Thus, we have 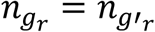 for all *r* in this simple case but not necessarily that 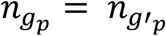. The four observations for genotype *g* may have different combinations of replicate number and batch number than the four observations from genotype *g’* and so the difference between the two genotypes will include differences due to replicates that end up in different batches. The amount of this effect will depend upon how many batch combinations contain both genotypes *g* and *g’*, which will also vary depending on the pair of genotypes considered and relative size of batch to the replicate. The estimator of 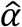 is no longer unbiased and contains differences due to batch:

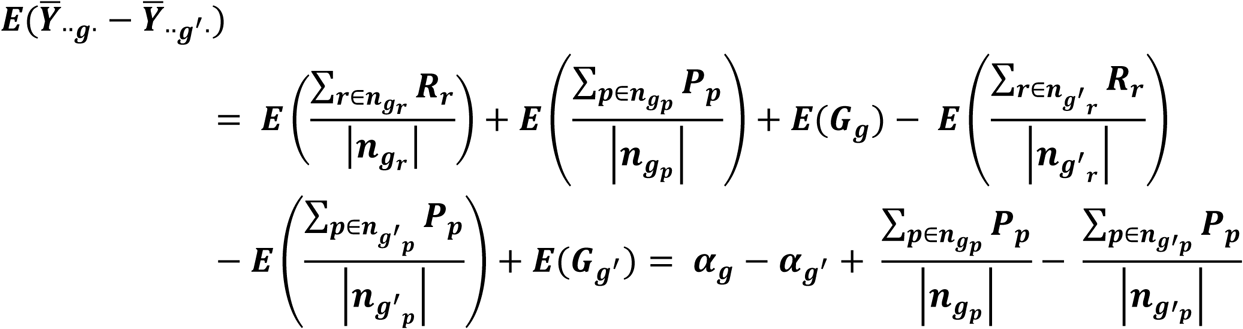

If all instances of *g* and *g’* are in the same batches then P = P’ and RR = CR.

#### Simulation 2: Conditional Randomization vs Re-Randomization for QTL effects in partial blocks

In a genetic reference panel experiment, it will not be uncommon to have more genotypes in one replicate than can fit into a single block. In addition to the sources of variation in *Simulation 1* there will be variance due to block. Further, it is often of interest to ask if the traits in one environment map to the same region as in another environment. For *f* = 1 to *f* blocks we drew the effect of each block from 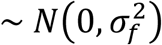. The number of blocks, *f* depends on the logistical constraints of sample collection and the number of genotypes to be assayed. Here we also consider batch sizes distinctly from block sizes. Batches have their own logistical constraints and these vary depending on the technology and the resources available. All other parameters are the same as in *Simulation 1*. In this particular simulation we simulated 4 reps * 384 genotypes = 1536 experimental units.

The trait value for an individual *i* in the incomplete block design is:

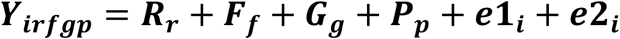

Following our notation, 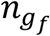 will denote the set of blocks for individuals having genotype g and now

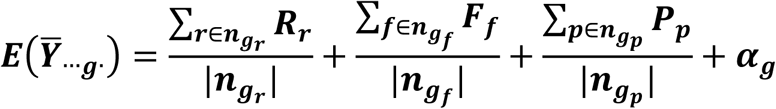

For simplicity, for both CR and RR we do not re-randomize the genotypes across replicates into batches. The replicate effect will cancel in the difference.

In CR, if genotype *g* and *g’* are in the same batch then by definition they were in the same block and the expected value for the difference between g and g’ is the genotypic difference as in Simulation 1. Across the population however, many of the comparisons between genotypes will be from different batches, and since we randomized genotypes into block locations, it is unlikely that two genotypes will consistently share a batch. The expected value of the difference between genotype *g* and *g’* will be

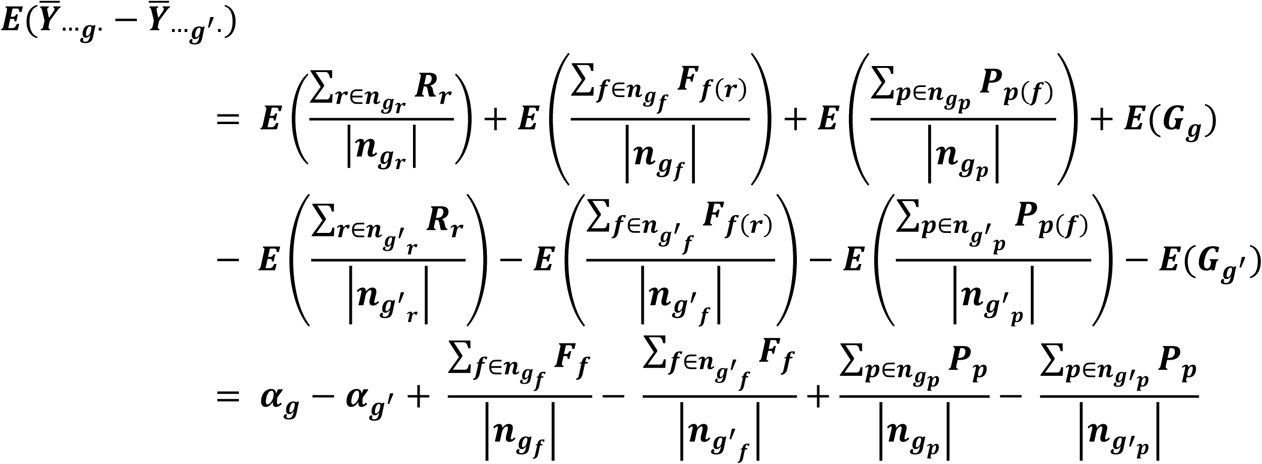

The maximum of the batch (block) effect will only occur when all the pairs of genotypes do not share a batch in any replicates. This is only a possibility if the batch size is equal to or smaller than the block size. If the batch size is larger than block size then the block and batch are no longer nested, and even if all genotypes are in different blocks, some blocks will be combined into the same batch and those batch effects will be shared and cancel out.

For RR, the experimental units are re-randomized from blocks into batches and the batch is not nested in the block. In RR if genotype *g* and *g’* are in the same batch then they may be in the same block, but they are likely to have been in different blocks. In CR, if the two genotypes shared a batch in any replicate then both the batch and block effect are eliminated in the expected difference. In RR that is no longer the case, and neither batch or block effects can be completely eliminated.

### Natural population simulations

#### Simulation 4. Conditional Randomization vs Re-Randomization across replicates

For g = 1 to 192 genotypes, the effect of genotypes was drawn from 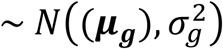 where genotypes have equal variance 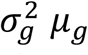. The means ***μ***_***g***_ were set such that *μ*_1_ =1 and *μ*_*g*+1_ -*μ*_*g*_=1. Other parameters are as in *Simulation 1*.

#### Simulation 5: *Conditional Randomization vs Re-Randomization for genotype and treatment effects*

Here we consider two scenarios an incomplete block and a split–plot.

*Incomplete Block:* We considered a simple two condition/treatment scenario, *t* = 1, 2 drawn from 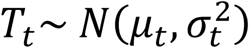 where treatments have equal variance 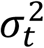 and different treatment means, *μ*_*t*_. Data were simulated for 192 genotypes as in *Simulation 4.*

In this particular simulation we simulated 4 reps * 2 treatments *192 genotypes = 1536 experimental units. For each replicate there were 8 blocks containing 24 genotypes (both treatments) for a total of 32 blocks in the experiment. All other parameters were simulated as in *Simulation 2*.

The trait value for an individual *i* was calculated as a sum of each of the sources of variation

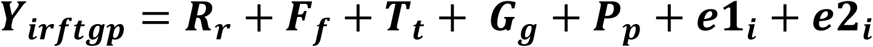

Now, let *n*_*t*_ be the set of individuals having treatment t, and |***n***_***t***_| be the number of individuals in *n*_*t*_, similarly 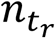 and 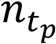 will denote the set of replicates and batches for individuals having treatment t. The expected mean trait value for treatment t is (without loss of generality, the expectation is conditional on the random effects rep, block, and batch):

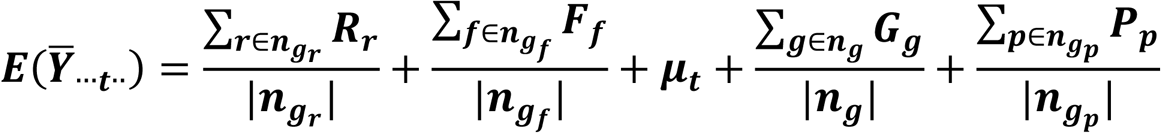

The generalized least squares estimator of the treatment effect between treatment t and t’ is:

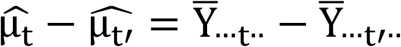

After sampling, experimental units were either kept together on a plate for data acquisition (CR) or randomized within the replicate after pairing the treatment pairs for the same genotype (P-RR). The estimated treatment effect is:

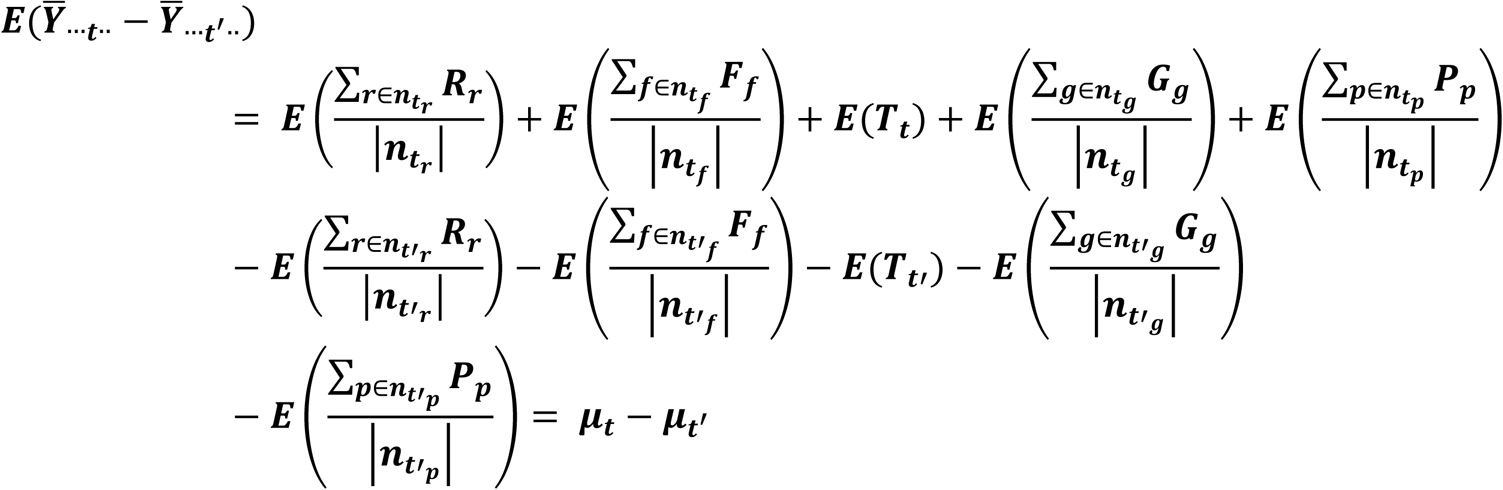

Since the re-randomization keeps the two treatments for the same genotype together here the expectation for the test of the treatment effect is the same in the CR and P-RR scenarios.

Spit plot: Some treatments cannot be implemented in the same block. As a result, experiment requires separate blocking typically referred to as a split-plot (SPD). The same data simulated in *Simulation 5* was used. However, individuals were assigned block and batch errors according to the SPD.

The replicate and block are different between treatments and CR will keep treatments in separate batches

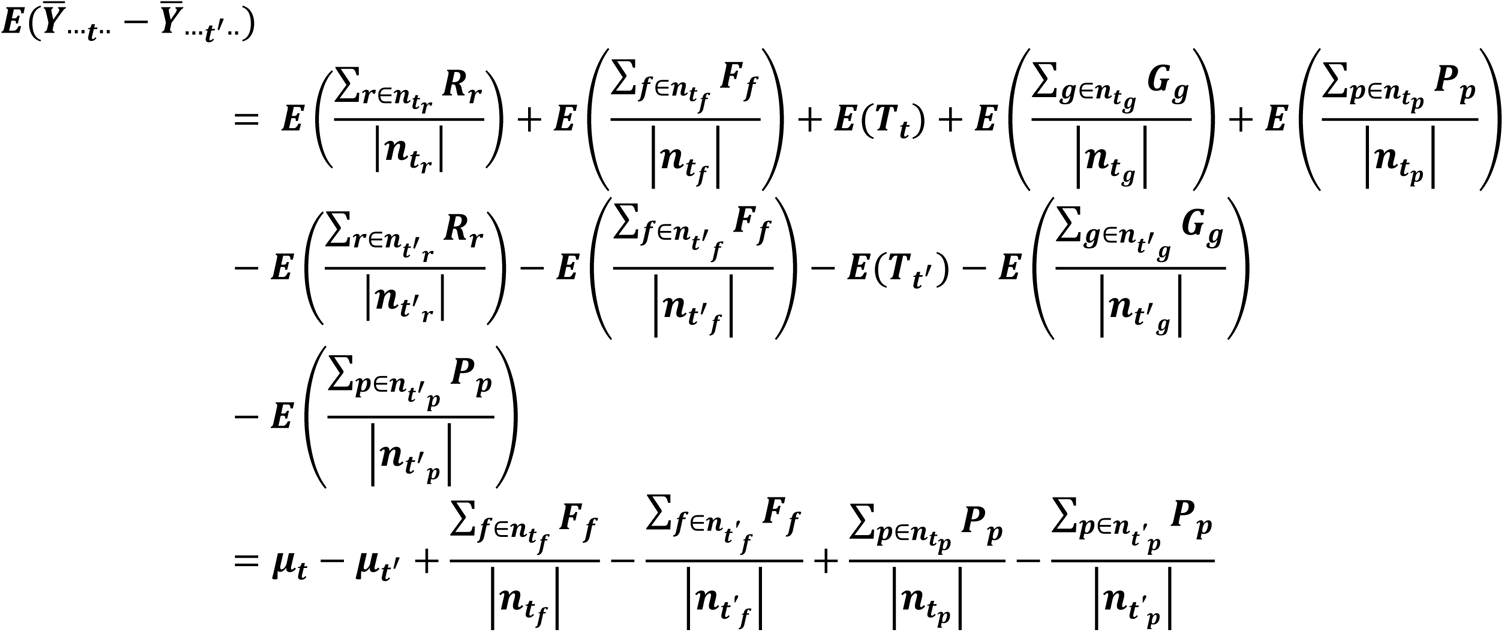

While in RR experimental units were randomized within the replicate keeping the treatment pairs for the same genotype from different blocks together in the same batch (P-RR).

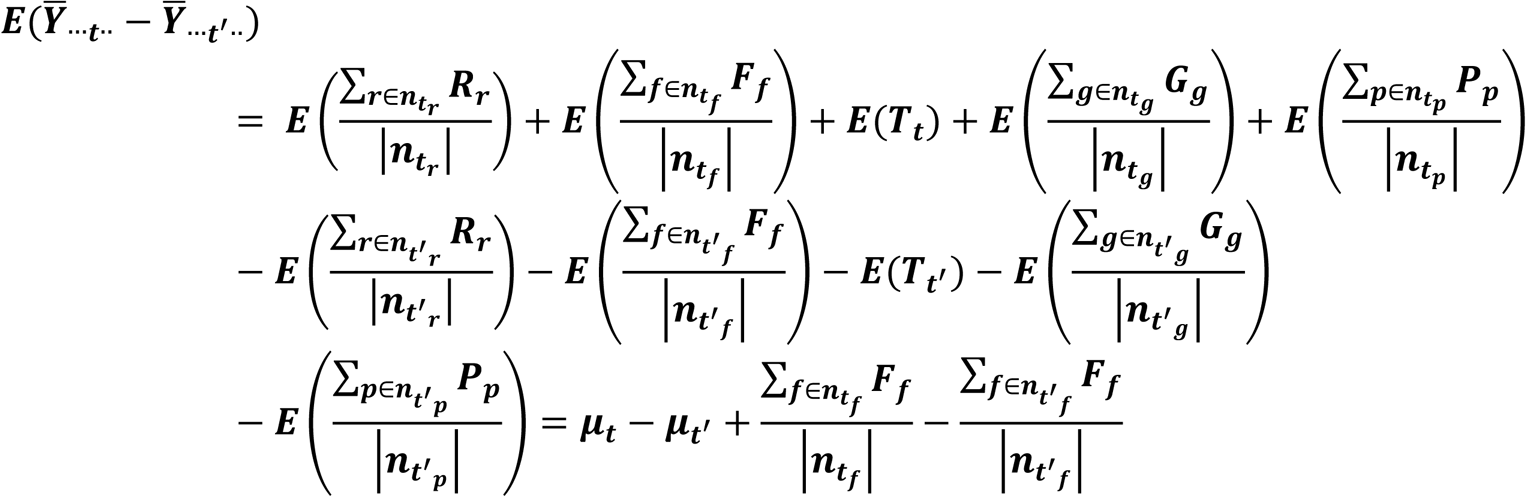

The IBD and SBD simulation conditions for to test RR and CR for power test of genotype and treatment. All combinations below were simulated for a total of 300 conditions with 50 iterations each.

---

**Supplementary Fig 1.**
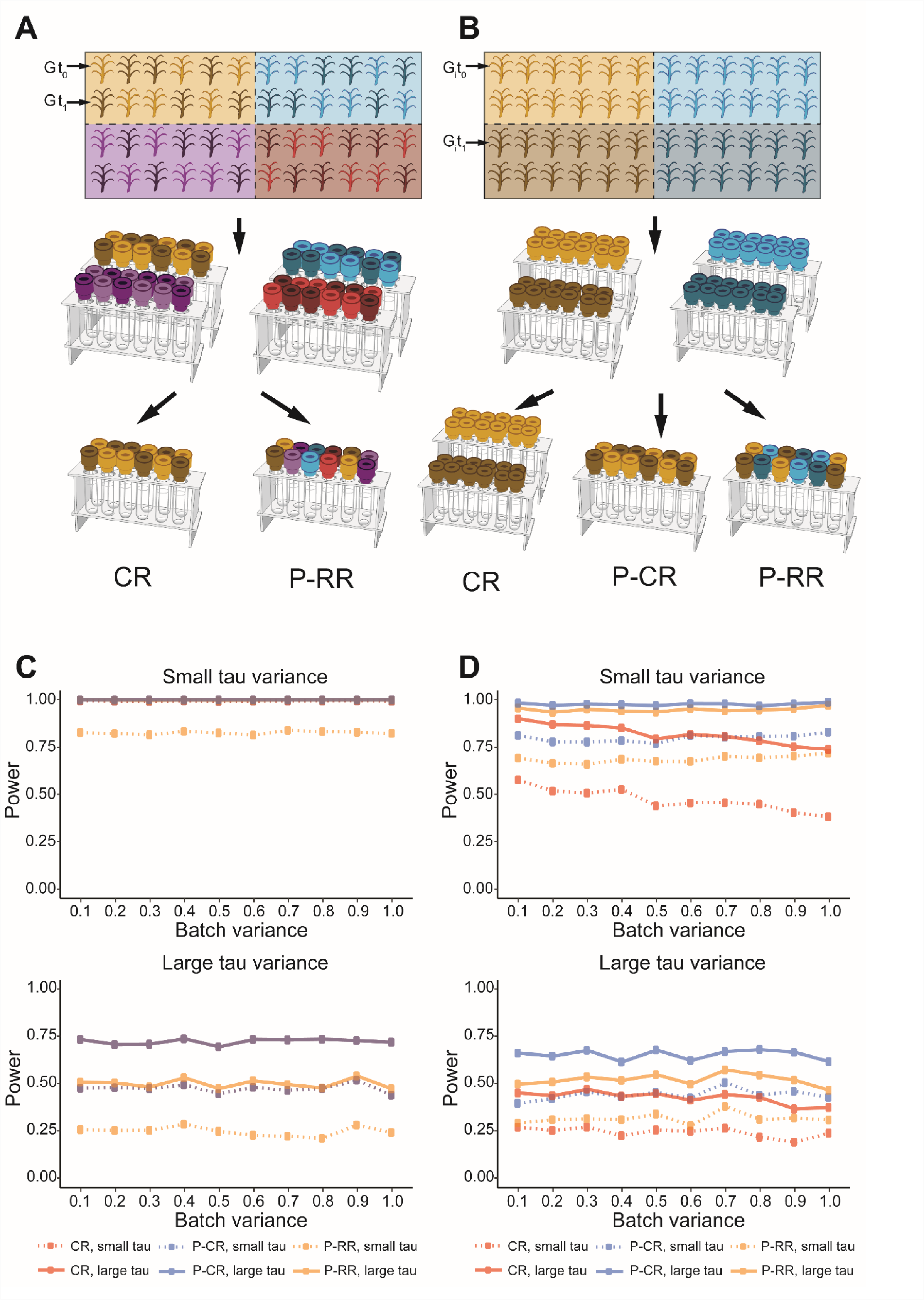
Incomplete block design vs split-plot design. (**A)** Design for sample collection and data acquisition in an incomplete block design (IBD) where all the treatments for a single genotype are co-located in the same block used for sampling. Samples can be assigned wither keeping the block structure (CR) or re-randomizing (RR); see also Figure 2 in the main manuscript. (**B)** A split plot (SPD) where treatments for a single genotype are sampled in different blocks. Samples can be assigned to batches either CR where treatment is then confounded with batch, or treatment sets can be completely re-randomized across blocks P-RR, or treatment sets can be combined and the block structure partially retained in the batches P-CR. (**C)** IBD design RR has less power than the CR regardless of treatment variance (low: variance = 0.5; high: variance = 2). In this comparison the treatment effects are paired at both sample collection and for data acquisition in CR while treatments are randomized in the block and again in the batch, some pairing will happen by chance. Note that the CR (red lines) and paired-CR (P-CR; blue lines) designs have the same result. (**D)** For SPD, treatments are in different blocks, keeping treatments separated (confounding treatment with batch; CR design) has much less power than pairing treatments of the same genotype (P-RR and P-CR designs; see also **Fig. 5**).

**Supplementary Fig 2.**
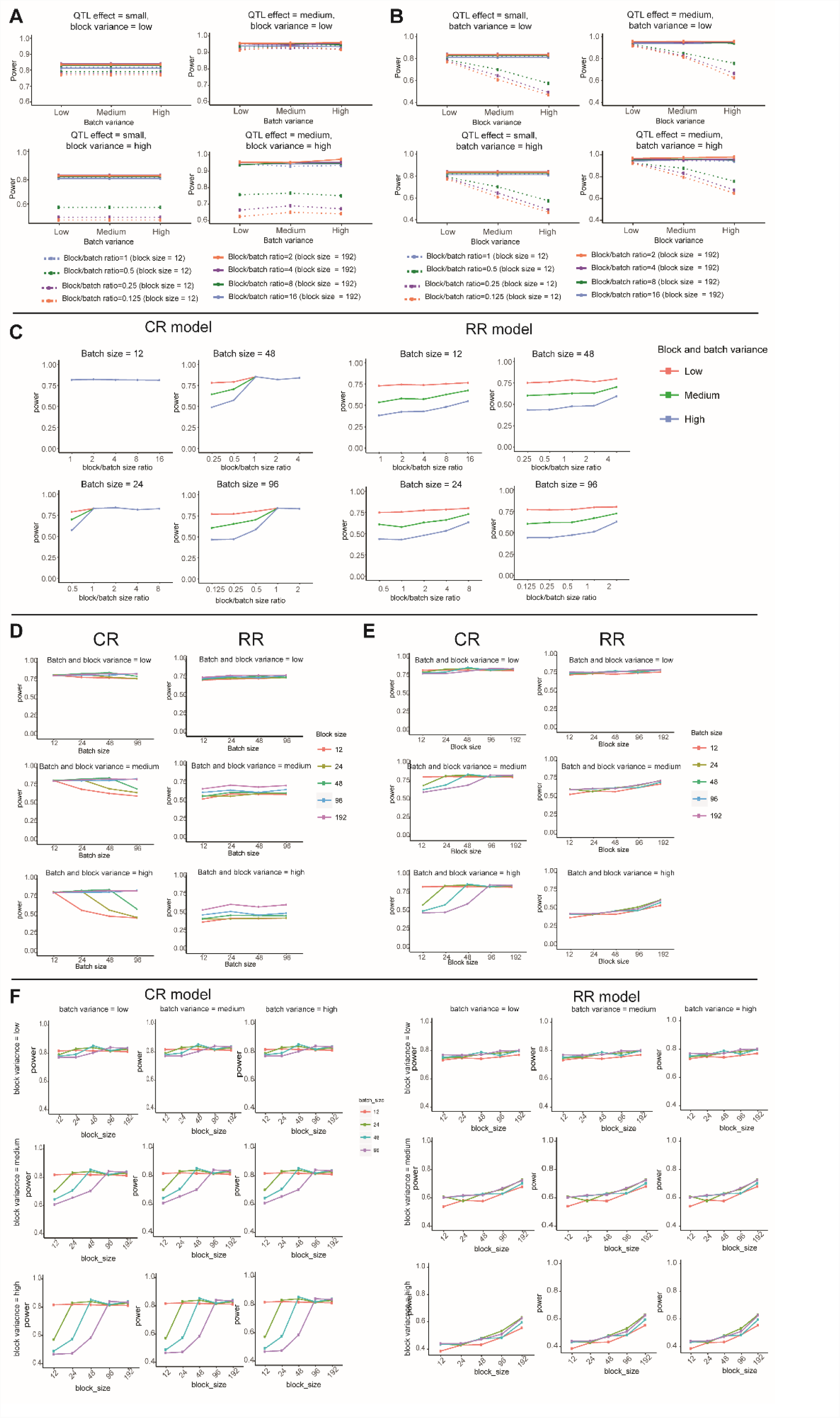
Conditional Randomization versus Re-Randomization for tests of genotypic effect. For all panels power is on the Y axis calculated as described in Figure 3 in the main manuscript. Effect of block and batch size on power. A simulation was conducted for a QTL with a single main effect for 192 RILs and 4 reps, with a QTL effect of 1.2 (small) and 1.5 (medium), replicate variance of 32 and error variance of 15. (A) When block variance is high (variance = 24), the power is higher when block/batch sizes are smaller (block variance = 24) and batch variance has little impact on power. When the block size is small, there is a loss of power when the variance is high, regardless of the size of the QTL effect. (B) When block variance is low block size does not matter. (C) Effect of the relative batch size to block size on power, when samples are CR or RR and batch and block variances are equal. In CR, batch variance is important only when batches are larger than blocks. (D) Effect of batch size and variance on power, when samples are CR or RR and batch and block variances are equal. In CR, there is a loss of power as variance increases and batch size is larger than block size. In RR, only the variance and block size impact power. (E) Effect of block size and variance on power, when samples are CR or RR and batch and block variances are equal. In CR, power increases as the block size increases relative to the batch size. In RR, larger block sizes have more power irrespective of batch size, although variance has a large influence on power. (F) Effect of batch and block size on power when batch and block variance are high and low, with QTL effect of 1.2, replicate variance of 32 and error variance of 15.

**Supplementary Fig 3.**
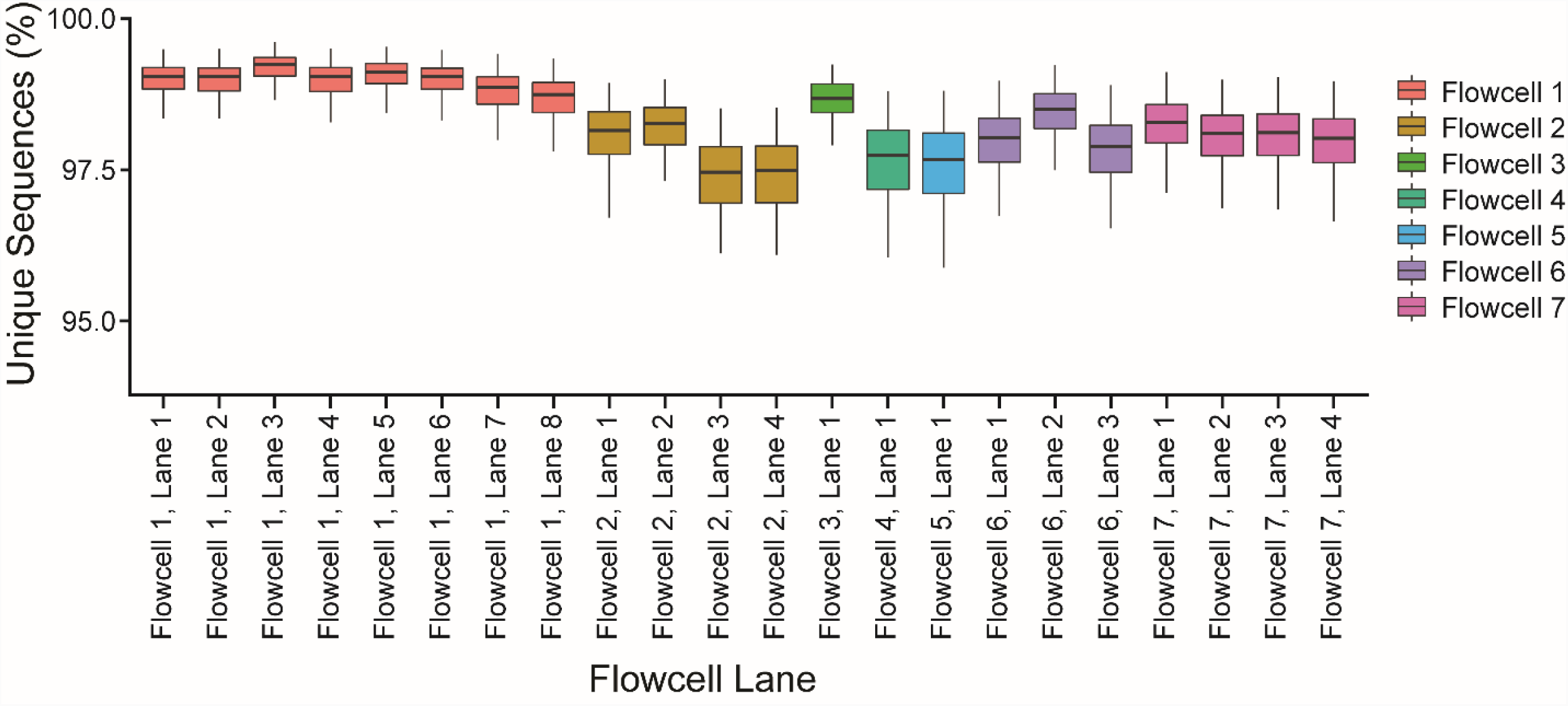
Flowcell lane effects on RNA-seq library complexity. A single multiplexed pool of a sample of 32 type 1 diabetes cases (3 cell types, 96 samples total; dbGaP study accession # phs001426.v1.p1, sequencing pool 1 samples; (1)). The known impact of lane (2) was the reason why the sample samples were run over multiple lanes and the data combined.

**Supplementary Fig 4.**
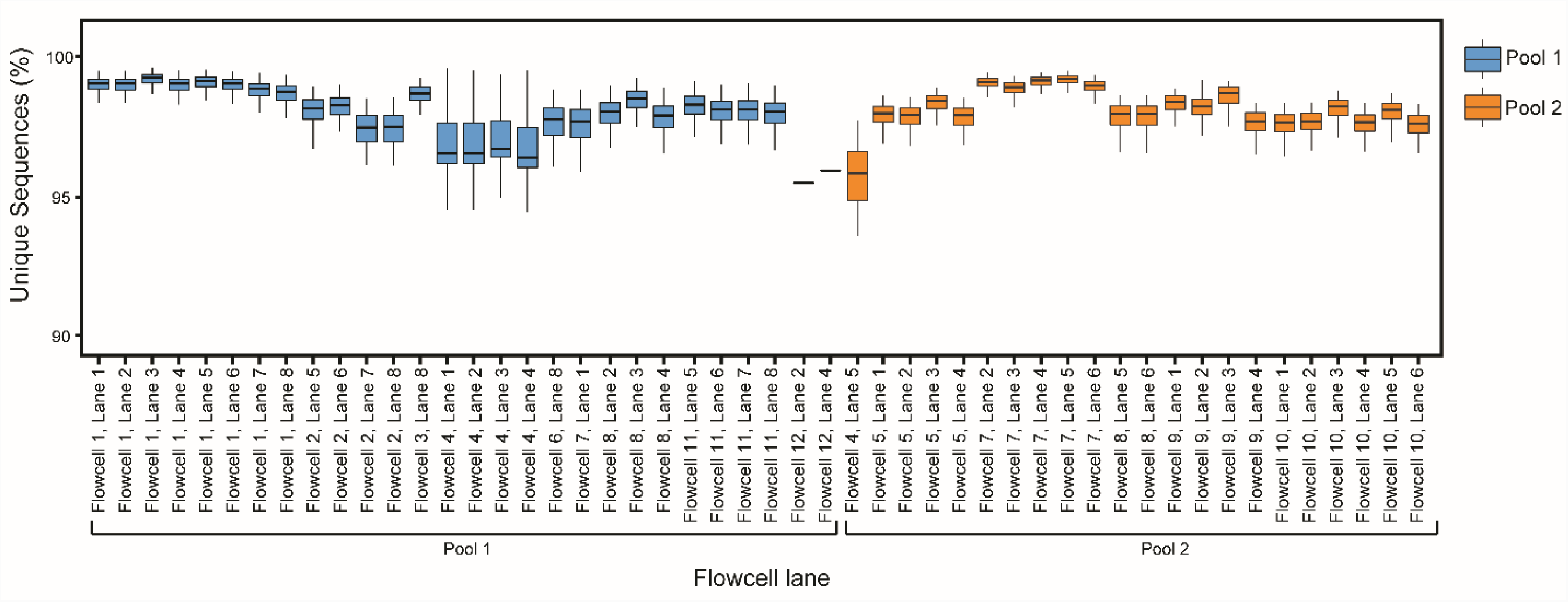
Flowcell lane effects on RNA-seq library complexity for two pools. Two multiplexed pools of 32 type 1 diabetes cases (3 cell types, 96 samples total; dbGaP study accession # phs001426.v1.p1, sequencing pool 1 and pool 2 samples; (1)). The pool effect was included in models of differential gene expression.

